# Handling and Adequacy of Iodine at Household Level: Community Based Cross-sectional Survey in Dega Damot District, West Gojjam Zone, Amhara Regional State, Ethiopia

**DOI:** 10.1101/586677

**Authors:** Aschalew Afework, Wondemagegn Mulu, Almayehu Abate, Abel Lule

## Abstract

**Introduction:** Iodine is essential for regulation of physical growth and neural development. Although, fortification of iodine has been practiced decades before and iodized salt is available, handling and cultural food preparation may affect the content of iodine in the dishes. Moreover, Dega Damot is mountainous area that may lose its iodine via erosion. Therefore, this study aimed at determining the handling and adequacy of iodine in the salt in Dega Damot district, West Gojjam Zone, Ethiopia.

**Method:** Community based cross-sectional study was conducted from October 24 to November 15, 2017 on 802 households. Stratified multi-stage sampling was employed to select households. A total of 422 and 380 house-holds from low land and high land, respectively took part in the study. For the interview either the husband or the spouse was selected randomly. Data on handling were collected by face-to-face interview using structured questionnaires. The concentration of iodine was determined using Rapid Test Kit. Descriptive statistics were used to describe relevant findings on the handling of iodized salt. Both bivariate and multivariate logistic regressions were analyzed to identify associated factors.

**Result:** From 802 samples tested, 37 (4.6%) had iodine greater than 15parts per million. The majority (94.5%) of the respondents have been roasting their salt for ‘Dikus’ preparation where as 91.1% of the households stored their salt in open containers. Salts with closed containers (AOR=3.22, CI=1.31-7.89), unroasted salts [AOR=5.23, CI=1.25-22.11], good knowledge on handling [AOR=5.55, CI=1.64-18.77], salts from high land area [AOR=2.11, 9CI=1.02-4.37], were significantly associated with adequacy of iodine Conclusions: Adequate utilization of iodine was very low. Roasting of salt was common. These phenomena may be continued because ‘Dikus’ preparation is cultured in this population. We recommend the supplementation of packed iodized salt in the dishes

## INTRODUCTION

Iodine is needed for regulation of physical growth and neural development. Iodine deficiency is the major cause of preventable mental retardation and its severity can range from mild mental blunting to cretinism [1].

Ethiopian public health institute 2016 survey reported that the national iodized salt utilization coverage was 89.2%. However, only about 26% of the surveyed households had salt that was adequately iodized (at ≥15 ppm). According to this report, the highest coverage of adequately iodized salt was in Tigray (55.2%) and Somali (49.4%) regions, and the lowest in the regions of Gambela (9.5%), SNNPR (13.7%), and Amhara (15%) [2]. The findings indicated that there is good national iodized salt coverage even if very low adequate iodized salt utilization at household level. This might be due to improper handling of the iodized salt. According to World Health Organization (WHO) and International Council for Control of Iodine Deficiency Disorders (ICCIDD) standard, elimination of IDD will be possible if more than 90% of the households consume adequately iodized salt [3].

Although Ethiopia, had set a national goal to eliminate IDD virtually by the year 2013 through universal salt iodization (USI) and increase access to iodized salt among households up to 90% [4]. The 2016 Ethiopian public health survey reported [3], only 26% adequate iodine coverage nationally. Moreover, there is great difference between the adequacy of iodine consumption at house hold level in Ethiopia and other study reports in different parts of the world. For example, 54.3% and 75.6% adequate iodine consumption at house hold level reported in Pakistan [5] and Ghana, respectively [6].

Factors such as duration of storage, size of the crystals, impurities, and the ambient temperature of the storage, humidity, cooking methods, time of adding and sunlight exposure influence the stability of iodized salt at the household level include [7, 8]. But varieties of improper handling have been reported in different studies. For example, a study done in Ghana, reported that 11.5% of households who stored salt in containers without a lid [6]; and 3.5 % of the respondents in Gondar town wash the salt to remove its impurities [9] and 64.9% of Assabi district add the salt during cooking [10].

Dega Damot district is one of the pocket areas of the country. Topographically, it consists of 35% mountainous, 30% ups and downs, 20% valleys and 15% plains. This topographic characteristic might be liable for erosion [11]. Thus, iodine may erode from the top area of the land. This characteristic of the soil with inadequate iodine content salt consumption may worsen the iodine deficiency. Even if high utilization of iodized salt in the country [2], improper handling might reduce the content of iodine in the dish and may not be sufficient to fulfill the demand of the body.

In Dega Damot district, there is cultural food preparation called ‘Dikus’. Dikus is prepared primary as raw material for the preparation of Wot (Ethiopian common cultural food). In this preparation, the iodized salt is either heated in sunlight or roasted in dry oven. This might lead to evaporation of iodine from the salt and might reduce the iodine content in the dish. Hence, the study determined the handling of salt and adequacy of iodine at house level in Dega Damot district, Ethiopia.

## Materials and Methods

### Study setting, design and period

A community based cross-sectional study was conducted in Dega Damot District from October 24-November 15, 2017. The District is located in West Gojjam Zone of Amhara National Regional State, 296 kilometers away from the capital city of Ethiopia. It is mountainous with high land and low land climate and produce different crops. The most commonly used foods in the high land are potato, barely and peens while lowland residents, commonly consumed maize, teff and barley foods. Erosion is very high in this area which may leads to the loss of iodine. According to the information obtained from the zonal finance and economy office based on the 2007 census, total population of the district is about 158, 667 with sex distribution of 78,115 males and 80,552 females. It has 32 sub-districts and the average number of households in each sub-district is estimated to be 830 with the average 6 members in each family [12]

### Sample size and Sampling procedure

The sample size was determined using single population formula with the following assumptions: level of confidence, 95%; margin of error, 5%; proportion of adequate iodine concentration from Lay Armachiho District, 29.7% [13]. By taking design effect of 2.5 and 5% contingency for the non–response, the sample size was 842. Stratified multi-stage sampling was used to select households from the source population. At first stage, 6 sub-districts (3 from high land and 3 from low land) were selected by lottery method. At the second stage, 6 villages (1 village from each district) were selected through lottery method. At the third stage, cluster sampling was used to select households from each village. Hence, 421 households from the low land and 421 from the high land were taken for this study. For the interview either the husband or the spouse was selected randomly.

### Data Collection

Demographic data and data for iodized salt handling were collected through face to face interview using structured questionnaires. The questionnaire contains open and close ended questions and included a section for observing the type of container used to store the salt and place of salt storage.

### Iodine determination

A teaspoon of salts collected from each household. The salts were tested for the content of iodine using the iodine rapid test kit (MBI Kits International). MBI KITS is improved iodized salt Field Test kit for salt fortified with potassium iodide. The interpretation of the result was based on color comparison with its chart (0, 1-15 and more than 15 parts per million, ppm). The cut-off proportion of 15 PPM and above was considered as adequately iodized salt using the WHO reference indicators for monitoring of iodized salt in the household level [14].

### Operational definitions

#### Adequate iodine in the salt

If the iodine content of the tested salt was > 15 parts per million

#### Low iodine content

If the iodine content of the tested salt was ≤ 15parts per million

#### No iodine

If the iodine content of the tested salt was 0 parts per million

#### Good knowledge on handling

If the respondent correctly answered greater than or equal to half of the knowledge questions items

#### Poor knowledge on handling

If the respondent correctly answered less than half of the knowledge questions item

### Data Analysis

The data was analyzed using SPSS version 20. Descriptive statistics such as frequencies and mean were used to describe the study participants in relation to relevant variables. The mean score of knowledge items questions was calculated to measure the level of knowledge of the respondents. Both bivariate and multivariate logistic regressions were used to identify associated factors. Odds ratio with 95% confidence interval were used to identify the presence and strength of association. .P-value less than 0.05 were taken as cut off point for statistical significance at the 95% confidence interval.

### Ethical Clearance

Ethical clearance was approved from Amhara Public health institute, and permission letter was taken from the district health office. After the purpose of the study was explained, verbal consent obtained from each respondent and participation was based on their willingness. Participants were assured that their names or personal identifiers are not included in the written questionnaire to ensure confidentiality. The findings were reported to the concerned bodies.

## Results

### Socio-demographic characteristics

A total of 802 households were took part in the study. Of them, 422 (52.6%) were from the low land. The mean ages of the respondents were 40-year-old (28-52). The average family size was six. Four hundred eighty two (60.1%) of the respondents were females and five hundred thirty four (66.2%) were unable to read and write. Details are depicted in table 1.

**Table1.**
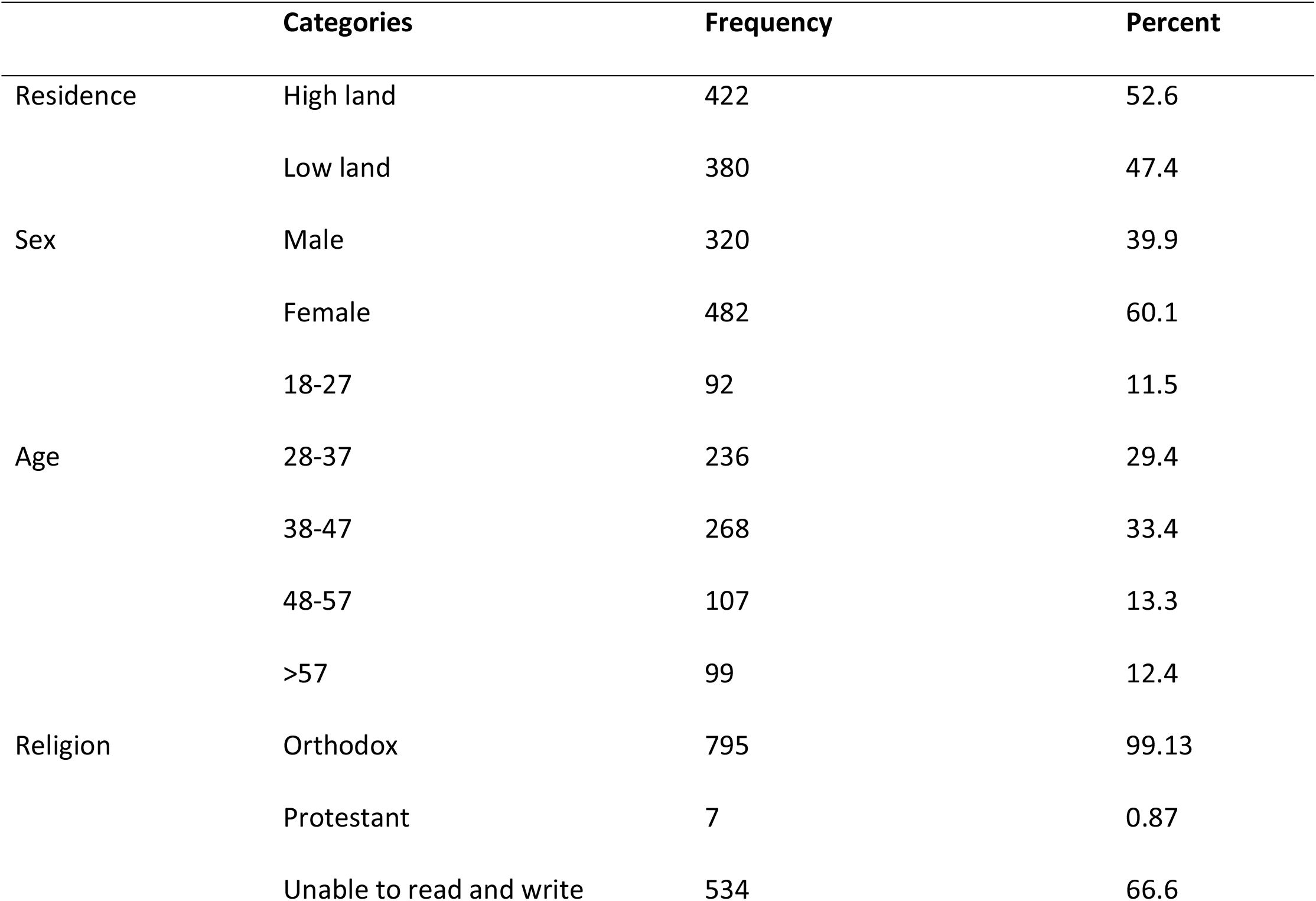
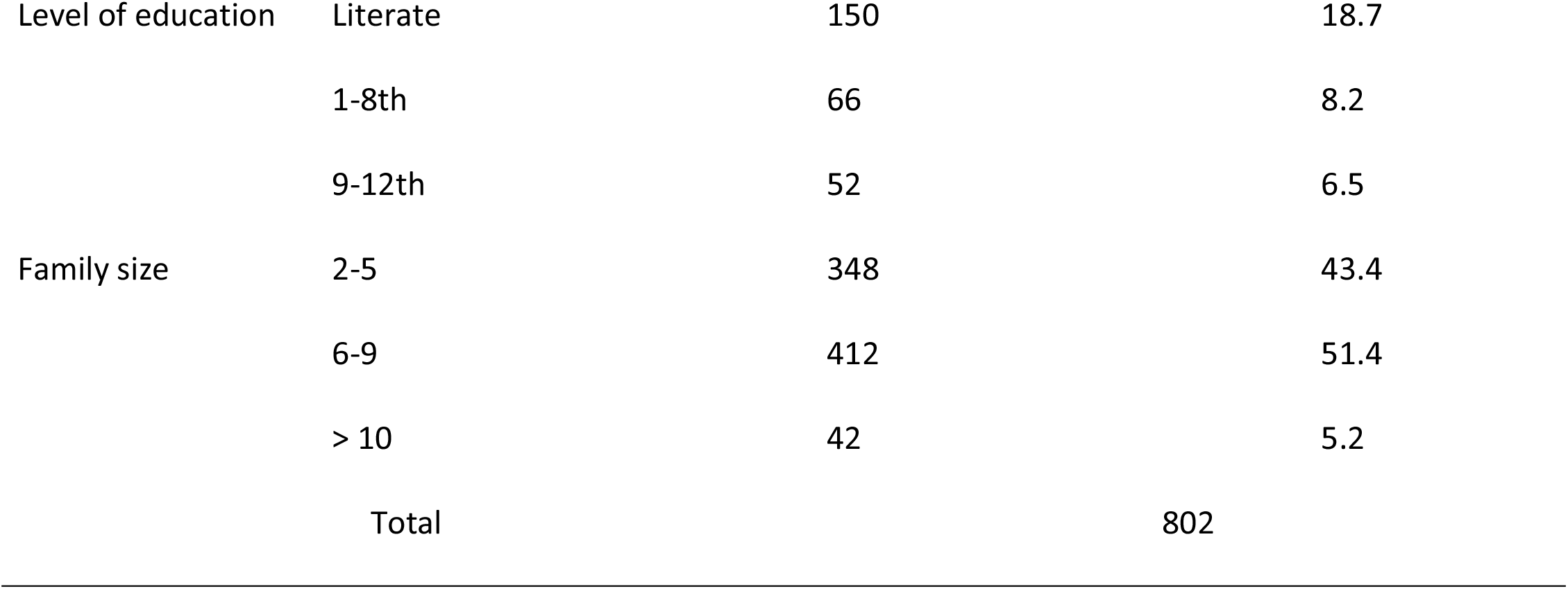
Demographic Characteristics of the respondents in Dega Damot District, October-2017.

### Handling of Iodized Salt

In this study, almost all (99.9) of the households had non-packed iodized salt. Of them, 85.8% were dried salts. Concerning frequency of buying, 534 (66.6%) of the households bought once a month and 96.5 percent of them buy the powdered one. The majority (91.1%) of the households stored their salt in an open container. Seven hundred twelve (88.8%) of them add the salt in the end stage of cooking. Most (94.5%) of the respondents had been roasting the salt while they prepare ‘Dikus’ (Table 2)

**Table2.**
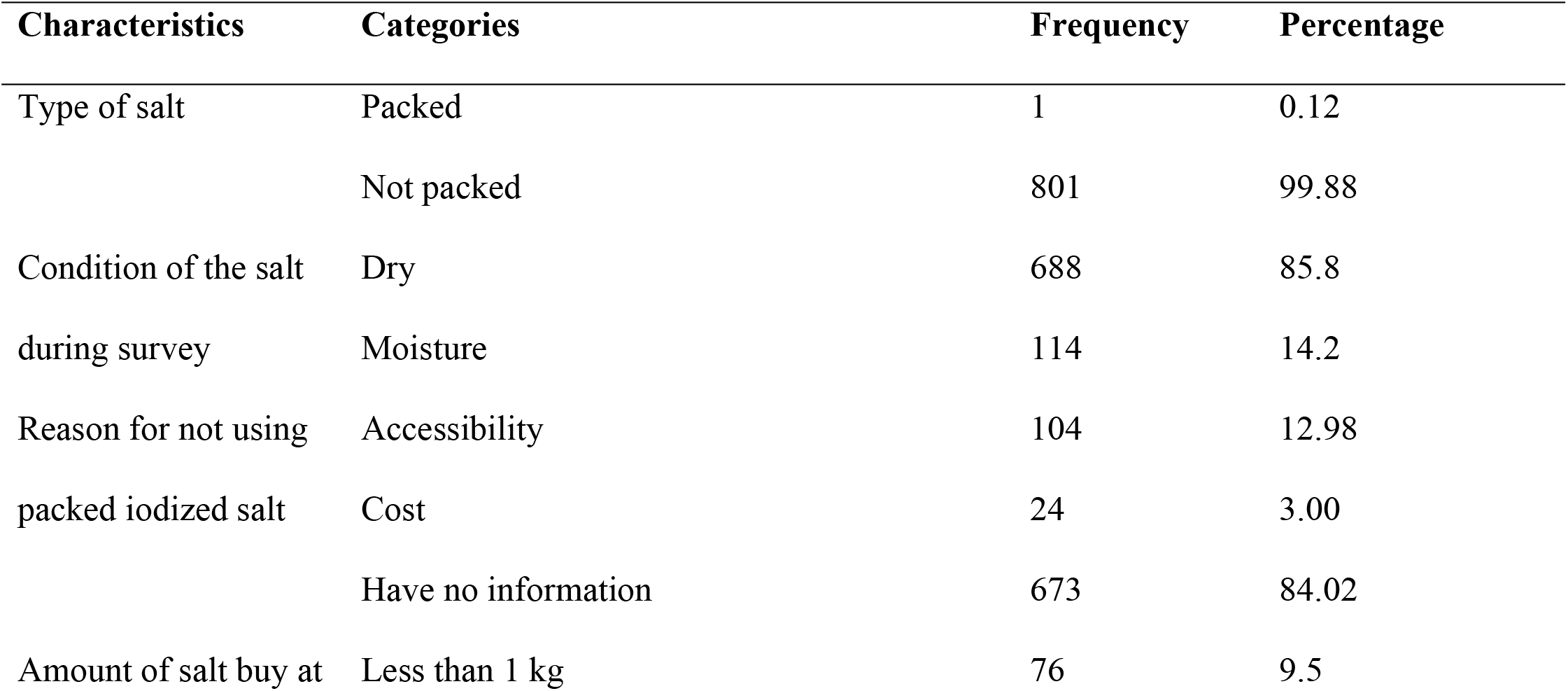
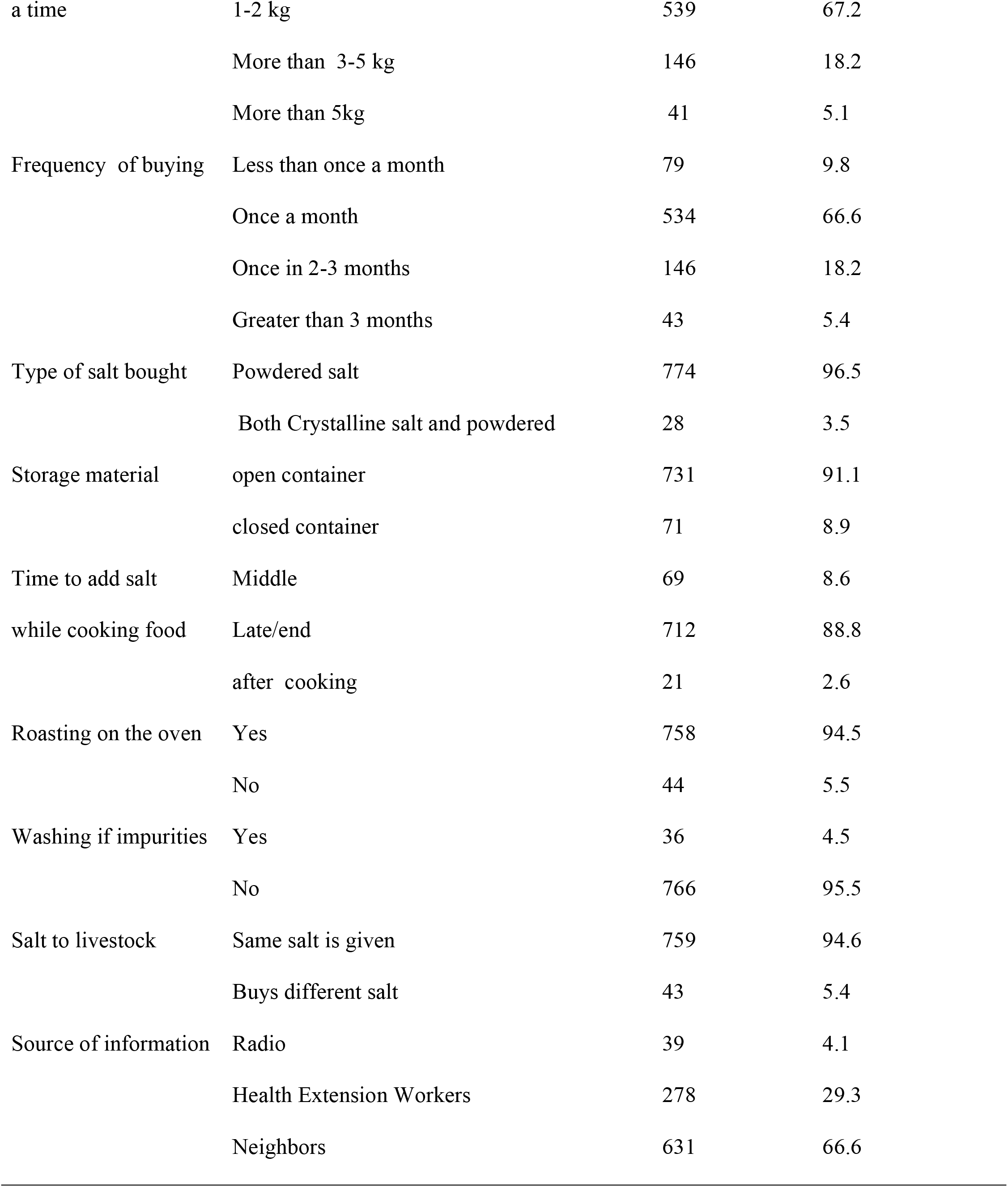
Handling and utilization of iodized salt in Dega Damot district.

| Characteristics | Categories | Frequency | Percentage |
| --- | --- | --- | --- |
| Type of salt | Packed | 1 | 0.12 |
|  | Not packed | 801 | 99.88 |
| Condition of the salt | Dry | 688 | 85.8 |
| during survey | Moisture | 114 | 14.2 |
| Reason for not using | Accessibility | 104 | 12.98 |
| packed iodized salt | Cost | 24 | 3.00 |
|  | Have no information | 673 | 84.02 |
| Amount of salt buy at | Less than 1 kg | 76 | 9.5 |
| a time | 1-2 kg | 539 | 67.2 |
|  | More than 3-5 kg | 146 | 18.2 |
|  | More than 5kg | 41 | 5.1 |
| Frequency of buying | Less than once a month | 79 | 9.8 |
|  | Once a month | 534 | 66.6 |
|  | Once in 2-3 months | 146 | 18.2 |
|  | Greater than 3 months | 43 | 5.4 |
| Type of salt bought | Powdered salt | 774 | 96.5 |
|  | Both Crystalline salt and powdered | 28 | 3.5 |
| Storage material | open container | 731 | 91.1 |
|  | closed container | 71 | 8.9 |
| Time to add salt | Middle | 69 | 8.6 |
| while cooking food | Late/end | 712 | 88.8 |
|  | after cooking | 21 | 2.6 |
| Roasting on the oven | Yes | 758 | 94.5 |
|  | No | 44 | 5.5 |
| Washing if impurities | Yes | 36 | 4.5 |
|  | No | 766 | 95.5 |
| Salt to livestock | Same salt is given | 759 | 94.6 |
|  | Buys different salt | 43 | 5.4 |
| Source of information | Radio | 39 | 4.1 |
|  | Health Extension Workers | 278 | 29.3 |
|  | Neighbors | 631 | 66.6 |

### Level of iodine concentration

From the total of 802 salts analyzed, 37 (4.6%) had adequate iodine content (greater than 15ppm) and 654 (81.6 %) had undetectable iodine content (Table 3)

**Table3.** Level of iodine concentration at household in Dega Damot District, West Gojjam, Ethiopia, October 2017.

| Variable | Frequency | Percent |
| --- | --- | --- |
| **Iodine content of the salt** |  |  |
| 0 parts per million | 654 | 81.6 |
| 1-15 parts per million | 111 | 13.8 |
| >15 parts per million | 37 | 4.6 |

### Multivariate analysis on the level of iodine

On multivariate analysis, salts in closed container [AOR=3.22; 95%CI=1.31, 7.89] have 3 times more likely to have concentration greater than 15ppm. Unroasted salts [AOR=5.23; 95%CI=1.25, 22.11] were positively associated with iodine concentration greater than 15ppm. Having good knowledge on handling [AOR=5.55; 95%CI=1.64, 18.77] significantly associated with iodine concentration greater than 15ppm. Salts collected from the high land area [AOR=2.11; 95%CI=1.02, 4.37] were 2 times more likely to have concentration greater than 15ppm. However, moisture content of the salt, educational status of the respondents, income, family size and frequency of buying were not associated with concentration of iodine salt in the salt (Table 4).

**Table4.** Multivariate analysis of factors that affect the iodine content of salt at Dega Damot District, October 2017.

| Variables | Iodine content |  | COR[95%CI] | AOR[95%CI] |
| --- | --- | --- | --- | --- |
|  | >15ppm | ≤15ppm |  |  |
| Storage Material |  |  |  |  |
| Closed container | 8 | 63 | 3.07 [1.35, 7.01] | 3.22 [1.31, 7.89] |
| Open container | 29 | 702 | 1.00 | 1.00 |
| Roasted iodized salt |  |  |  |  |
| No | 35 | 588 | 5.27 [1.26, 22.12] | 5.23 [1.25, 22.11] |
| Yes | 2 | 177 | 1.00 | 1.00 |
| Educational status of the respondents |  |  |  |  |
| No educated individuals | 28 | 506 | 0.71 [0.16, 3.06] | 0.67 [0.15, 3.01] |
| Elementary education | 7 | 208 | 1.17 [0.23, 5.78] | 1.06 [0.20, 5.73] |
| Secondary educated individuals | 2 | 51 | 1.00 | 1.00 |
| Respondents Knowledge on Handling |  |  |  |  |
| Poor | 32 | 742 | 1.00 | 1.00 |
| Good | 5 | 23 | 5.04 [1.80, 14.12] | 5.55 [1.64, 18. 77] |
| Income |  |  |  |  |
| Less than or equal to 11, 999 | 35 | 750 | 1.00 | 1.00 |
| More than 12, 000 | 2 | 15 | 2.86 [0.63, 12.98] | 1.69 [0.23, 12.57] |
| Salt collected site |  |  |  |  |
| Low land area | 13 | 409 | 1.00 | 1.00 |
| High land area | 24 | 356 | 2.12 [1.06, 4.23] | 2.11 [1.02, 4.37] |
| Condition of the salt during survey |  |  |  |  |
| Dry /normal | 29 | 669 | 1.00 | 1.00 |
| Moist salt | 8 | 96 | 1.92[0.85, 4.33] | 0.53 [0.23, 1.19] |
| **Frequency of bought** |  |  |  |  |
| Less than once a month | 4 | 75 | 1.00 | 1.00 |
| Once a month | 25 | 509 | 1.06 [0.37, 3.20] | 1.16 [0.49, 3.59] |
| Once in 2-3 months | 5 | 141 | 1.50 [0.39, 5.77] | 2.06 [0.49, 8.56] |
| Greater than 3 months | 3 | 40 | 0.71 [0.15, 3.33] | 0.73 [0.14, 3.82] |
| **Family size** |  |  |  |  |
| 2-5 | 18 | 330 | 0.45 [0.06, 3.44] | 0.43 [0.56, 3.34] |
| 6-9 | 18 | 394 | 0.534 [0.07, 4.10] | 0.52 [0.07, 3.99] |
| Greater than 9 | 1 | 41 | 1.00 | 1.00 |

## DISCUSSION

A community based cross-sectional study attempted to assess handling and adequacy of iodine in the salt at the household level. Proper handling of the iodized salt at household level is a critical point to achieve the intended intervention of iodine fortification. In this study, only 4.6% of households had adequate iodine content in their salts. This finding is very far from WHO recommendation to eliminate iodine deficiency (3) and also with the Ethiopian national plan to eliminate IDD virtually by the year 2013 through adequate iodized salt utilization (4). Moreover, the present finding is significantly lower than 2016 Ethiopia Public health survey report [2]. This is an alarm to work at the community level in order to achieve the intended WHO goals.

Comparing with others, our study finding is by far lower than in Belgaum district, Indian [15], Pakistan [5] and Ghana [6] on which 33.3%, 54.3% and 44.8% of the household had adequate iodized salt, respectively. Moreover, the proportion of adequate iodine consumption in this population was lower than studies done in Gondar [9] and Labo Assabi district [10]. The reason might be due to poor handling of the salt in the study area or cultural food preparation led to high loss of iodine from the salt.

In the present study, 94.5% of respondents reported roasting the salt while preparing Dikus (the raw material for the preparation of ‘Wot’ (common cultural food in Ethiopia)). This cultural food preparation led to the common phenomena of roasting of salt in this population. Roasting is not common in other studies even if exposure to sunlight is common practice [10, 17, 18]. This might be one of the reasons for the loss of iodine from the salt. Moreover, almost all (99.9%) of the households used non-packed iodized salt. The result is lower compared to studies done in Wolaita Sodo town [17] and Dera District, Ethiopia [18], the packed iodized salt utilization was 11.1% and 12.49%, respectively. High proportion of roasting of salts and lower utilization of packed iodized salt needs attention in this population.

In this study, the roasted salts were 5.2 times less likely to have adequate iodine than the counter parts. Even if it is not identical to that of roasting, exposure to sunlight in the other similar studies were also associated with inadequacy of iodine content. For example, not exposing salt to sunlight was positively associated with utilization of adequately iodized salt in LaboAssabi District [11]. In a similar previous study at Gondar, Ethiopia, those who did not expose salt to sunlight were 7.26 times more adequate iodized salt than those who exposed salt to sunlight [10]. Moreover, exposure of salt to sunlight were about 4 times more likely to have adequately iodized salt as compared to their counter parts in a similar study done in Wolaita Sodo town[16]

In this study, respondents who had good knowledge on handling salts were 5.6 times more likely to have iodine concentration greater than 15ppm.Comparatively, respondents who had good knowledge on handling salts were 1.94 times more likely to have adequate iodine than those who do have poor knowledge in a study conducted at Gondar town, Ethiopia [10]. Likewise, knowledge of the respondents was associated with adequacy of iodine content in the salt in Dera District, Ethiopia [17]. However, knowledge about iodized salt has no significant association with iodine content in LaboAssebi District, Ethiopia [11].

### Conclusion and recommendation

The proportion of households with adequate iodine salt is very low. Roasting of the salt is a common practice in this population. These phenomena may be continued in the future because ‘Dikus’ preparation is cultured in this population. We recommend the utilization of packed iodized salt in the dish to attain the deficit due to roasting of salts during Dikus preparation

## ACKNOWLEDGMENT

We are grateful for GAMBY Medical and Business College in providing conducive environment for the dissemination of the finding of this research. We would like to extend our appreciation to Amhara National Regional State Health Bureau, Research and Technology Transfer Core Process for its ethical approval. Our acknowledgment also goes to Dega Damot health office for its letter of permission. We acknowledge UNICEF, Bahir Dar branch for materials supporting for the chemical analysis. We also thank the respondents. Finally our deepest appreciation goes to the data collector: Tamirat Afework and Nibret Tilahun

